# Response preparation involves a release of intracortical inhibition in task-irrelevant muscles

**DOI:** 10.1101/2020.06.23.167965

**Authors:** Isaac N. Gomez, Kara Ormiston, Ian Greenhouse

**Author notes:** Corresponding Author Ian Greenhouse 181 Esslinger Hall 1525 University St. Eugene, OR 97403.

## Abstract

Action preparation involves widespread modulation of motor system excitability, but the precise mechanisms are unknown. In this study, we investigated whether intracortical inhibition changes in task-irrelevant muscle representations during action preparation. We used transcranial magnetic stimulation (TMS) combined with electromyography in healthy human adults to measure motor evoked potentials (MEPs) and cortical silent periods (CSPs) in task-irrelevant muscles during the preparatory period of simple delayed response tasks. In Experiment 1, participants responded with the left-index finger in one task condition and the right-index finger in another task condition, while MEPs and CSPs were measured from the contralateral non-responding and tonically contracted index finger. During Experiment 2, participants responded with the right pinky finger while MEPs and CSPs were measured from the tonically contracted left-index finger. In both experiments, MEPs and CSPs were compared between the task preparatory period and a resting intertrial baseline. The CSP duration during response preparation decreased from baseline in every case. A laterality difference was also observed in Experiment 1, with a greater CSP reduction during the preparation of left finger responses compared to right finger responses. MEP amplitudes showed no modulation during movement preparation in any of the three response conditions. These findings indicate cortical inhibition associated with task-irrelevant muscles is transiently released during action preparation and implicate a novel mechanism for the controlled and coordinated release of motor cortex inhibition.

**New & Noteworthy:** In this study we observed the first evidence of a release of intracortical inhibition in task-irrelevant muscle representations during response preparation. We applied transcranial magnetic stimulation to elicit cortical silent periods in task-irrelevant muscles during response preparation and observed a consistent decrease in the silent period duration relative to a resting baseline. These findings address the question of whether cortical mechanisms underlie widespread modulation in motor excitability during response preparation.

## Introduction

Successful goal-directed behavior depends on the ability to use information in the environment to prepare the motor system for action. Our current understanding of the neural mechanisms involved in action preparation in humans is incomplete. Several transcranial magnetic stimulation (TMS) studies have found evidence for inhibition of the motor output pathway during the preparation of actions, referred to as preparatory inhibition, measured as a reduction in motor evoked potential (MEP) amplitudes following an informative cue in a delayed response task (Davranche et al., 2007; Duque et al., 2010; Duque & Ivry, 2009; Greenhouse et al., 2015; Hannah et al., 2018; Hasbroucq et al., 1997; Ibáñez et al., 2020; Labruna et al., 2019; Lebon et al., 2016; Quoilin et al., 2016; Sinclair & Hammond, 2008; Vassiliadis et al., 2020). Further evidence suggests preparatory inhibition is widespread, influencing not only task-relevant muscles, but task-irrelevant muscles as well (Duque & Ivry, 2009; Greenhouse et al., 2015). Whether this widespread decrease in motor system excitability during action preparation reflects the influence of local intracortical, long-distance transcortical, or subcortical mechanisms is unclear.

The cortical silent period (CSP) is the suppressed activity in the electromyogram (EMG) of a tonically active muscle following a single TMS pulse (Wilson et al., 1993). Early and late portions of the CSP reflect spinal and cortical inhibitory mechanisms, respectively (Chen et al., 1999; Fuhr et al., 1991). Pharmacological studies suggest a mixture of GABA_A_ and GABA_B_ mediated intracortical inhibitory mechanisms contribute to the latter portion of the CSP (Inghilleri et al., 1996; McDonnell et al., 2006; Paulus et al., 2008; Werhahn et al., 1999).

During action preparation, the duration of the CSP measured from a muscle involved in the planned response progressively shortens over the course of the fore-period - likely the result of increasing neural drive approaching movement onset (Davranche et al., 2007). This release of inhibition, seen in the CSP, occurs in parallel with a decrease in MEP amplitude (Davranche et al., 2007). The collective existing evidence suggests a local release of intracortical inhibition occurs in the context of a more widespread decrease in corticospinal excitability, possibly via a subcortical or inter-cortical mechanism. However, while previous CSP studies have focused on task-involved muscles, to our knowledge, no previous work has investigated the CSP in task-irrelevant muscles, and the question of whether changes in intracortical inhibition are specific to task-involved muscles remains unanswered. Furthermore, investigating the CSP in task-irrelevant muscles avoids issues related to changes in neural drive associated with response execution.

In the current study, we sought to assess whether intracortical inhibitory mechanisms contribute to the widespread modulation of motor system excitability during action preparation. To this end, we examined MEP amplitudes and CSP durations in a task-irrelevant muscle during action preparation in a delayed response task. We tested two competing hypotheses: 1) CSP duration would be shorter during preparation when compared to a resting baseline, reflecting a widespread release of intracortical inhibition during action preparation. Such a pattern would parallel the pattern observed in a muscle selected for a forthcoming response, implicating a common mechanism. 2) Alternatively, CSP duration would be longer during preparation when compared to baseline. Such a pattern would be consistent with the recruitment of widespread intracortical inhibition during action preparation, implicating a specific cortical mechanism for preparatory inhibition.

## Materials and Methods

### Participants

A total of 22 healthy, right-handed participants (10 females, 12 males, 24 ± 5 years of age) were included in the study. Data were collected in two experiments (n = 14 in each). Six participants completed both experiments, and 8 participants were unique to each experiment. Participant characteristics are presented in Table 1. All participants were screened for contraindications to TMS and provided written informed consent per a protocol approved by the IRB of the University of Oregon.

**Table 1.** Participant Characteristics

|  | Participant | Sex | Age | Left FDI MVC | Right M1 RMT | Right FDI MVC | Left M1 RMT |
| --- | --- | --- | --- | --- | --- | --- | --- |
| **Experiment 1** | 1 | m | 26 | 7.41 | 44 | 6.96 | 48 |
|  | 2 | m | 22 | 9.91 | 62 | 8.74 | 58 |
|  | 3 | m | 22 | 4.86 | 42 | 9.91 | 42 |
|  | 4 | f | 20 | 3.32 | 54 | 2.06 | 54 |
|  | 5 | m | 32 | 5.94 | 48 | 7.06 | 54 |
|  | 6 | f | 20 | 3.84 | 58 | 3.84 | 52 |
|  | 7 | f | 21 | 9.91 | 47 | 8.79 | 44 |
|  | 8 | m | 29 | 7.62 | 50 | 5.06 | 48 |
|  | 9 | f | 19 | 9.91 | 36 | 9.91 | 36 |
|  | 10 | m | 22 | 4.53 | 41 | 4.12 | 40 |
|  | 11 | m | 28 | 8.05 | 36 | 9.91 | 48 |
|  | 12 | m | 20 | 9.91 | 54 | 9.91 | 51 |
|  | 13 | f | 26 | 8.92 | 48 | 6.90 | 44 |
|  | 14 | m | 32 | 8.36 | 47 | 8.87 | 39 |
| **Experiment 2** | 1 | m | 26 | 5.95 | 44 |  |  |
|  | 2 | m | 22 | 9.91 | 52 |  |  |
|  | 3 | m | 22 | 6.04 | 41 |  |  |
|  | 4 | f | 20 | 2.66 | 55 |  |  |
|  | 5 | m | 32 | 6.18 | 48 |  |  |
|  | 6 | f | 20 | 3.84 | 58 |  |  |
|  | 15 | f | 19 | 1.49 | 36 |  |  |
|  | 16 | f | 21 | 1.86 | 34 |  |  |
|  | 17 | m | 19 | 2.39 | 64 |  |  |
|  | 18 | f | 19 | 2.78 | 43 |  |  |
|  | 19 | m | 19 | 3.74 | 35 |  |  |
|  | 20 | f | 19 | 6.53 | 45 |  |  |
|  | 21 | f | 30 | 6.89 | 40 |  |  |
|  | 22 | m | 25 | 6.99 | 46 |  |  |

MVC = maximum voluntary contraction (mV), RMT = resting motor threshold (percent maximum stimulator output). Participants 1 through 6 completed both experimental protocols.

### Experimental Setup

Participants were seated comfortably in front of a computer monitor with both hands placed palm-down on the surface of a table. USB-interfaced response buttons were fixed to a button-box such that button presses could be executed starting from a resting hand position. The configuration of the response buttons differed between Experiments 1 and 2 (see below). Visual stimulus presentation was controlled by Psychtoolbox 3.0, and both EMG recording and the timed administration of TMS pulses were controlled by the VETA toolbox (Jackson & Greenhouse, 2019) in Matlab. All experimental task code, analysis code, and data are available for download through the Open Science Framework at https://osf.io/jvnmq/.

### Transcranial Magnetic Stimulation & Electromyography

Surface EMG was recorded using bipolar electrodes placed over both FDI muscles for Experiment 1, and the left FDI and right ADM in Experiment 2. A ground electrode was placed over the ulnar styloid process of the left arm. EMG was sampled at 5,000 Hz, amplified 1000 x, and bandpass filtered (50-450 Hz; Delsys). At the start of the experiment, maximum voluntary contraction (MVC) of the target FDI muscle was determined using a foam squeeze ball placed between the left-index and thumb finger. Participants executed four consecutive 4 s contractions separated by 1 s of rest, and the maximum peak-to-peak amplitude of the EMG activity was calculated. Subsequently, participants were trained in maintaining a tonic contraction of near 25% MVC while holding the foam squeeze ball and visualizing the live EMG trace with markers indicating the target amplitude. In Experiment 1, the determination of MVC was done separately for the left and right FDI corresponding to the two response conditions. In Experiment 2, MVC was determined for the left FDI only.

TMS was administered using a Magstim 200-2 stimulator with a 7-cm-diameter figure-of-eight coil. The center of the TMS coil was positioned over the left M1 to elicit MEPs and CSPs in the right FDI muscle (Experiment 1 only), and over the right M1 to elicit MEPs and CSPs in the left FDI muscle (Experiment 1 and 2). A standard hotspotting and thresholding procedure was used while the participant remained at rest. First, the coil was positioned approximately 2 cm anterior and 5 cm lateral to the vertex, over the hemisphere contralateral to the target muscle, and with the coil oriented approximately 45 degrees off the midline to induce a current in the posterior to anterior direction. Second, the TMS intensity was adjusted and the coil was repositioned in incremental adjustments of ~1 cm until consistent MEPs were elicited from the targeted FDI. During this hotspotting procedure, TMS pulses were administered once every 4 s. Third, once the optimal coil position and orientation were determined, a felt-tip marker was used to trace the coil position directly on the participant’s scalp. Finally, the resting motor threshold (RMT) was determined as the intensity of TMS, which elicited MEPs with amplitudes of at least 50 μV on 5 out of 10 attempts. During subsequent testing, TMS was administered at 115% RMT. The average RMTs for Experiments 1 and 2 were 47 ± 7% and 46 ± 9% of maximum stimulator output, respectively.

### Unimanual Delayed Response Task

Participants completed a unimanual delayed response task while maintaining a tonic contraction with the non-responding hand (Figure 1A and B). Each trial of the task consisted of a 200 ms baseline fixation cue, followed by a 900 ms preparatory cue and a 500 ms imperative Go stimulus (Figure 1C). Each block consisted of 44 Go trials and six randomly interspersed catch trials, in which the preparatory cue remained on the screen through the end of the trial, and the Go stimulus never appeared. Catch trials were included to discourage premature responses. Participants were instructed to keep the responding hand at rest between trials and to respond as quickly as possible to the Go stimuli.

**Figure 1.**
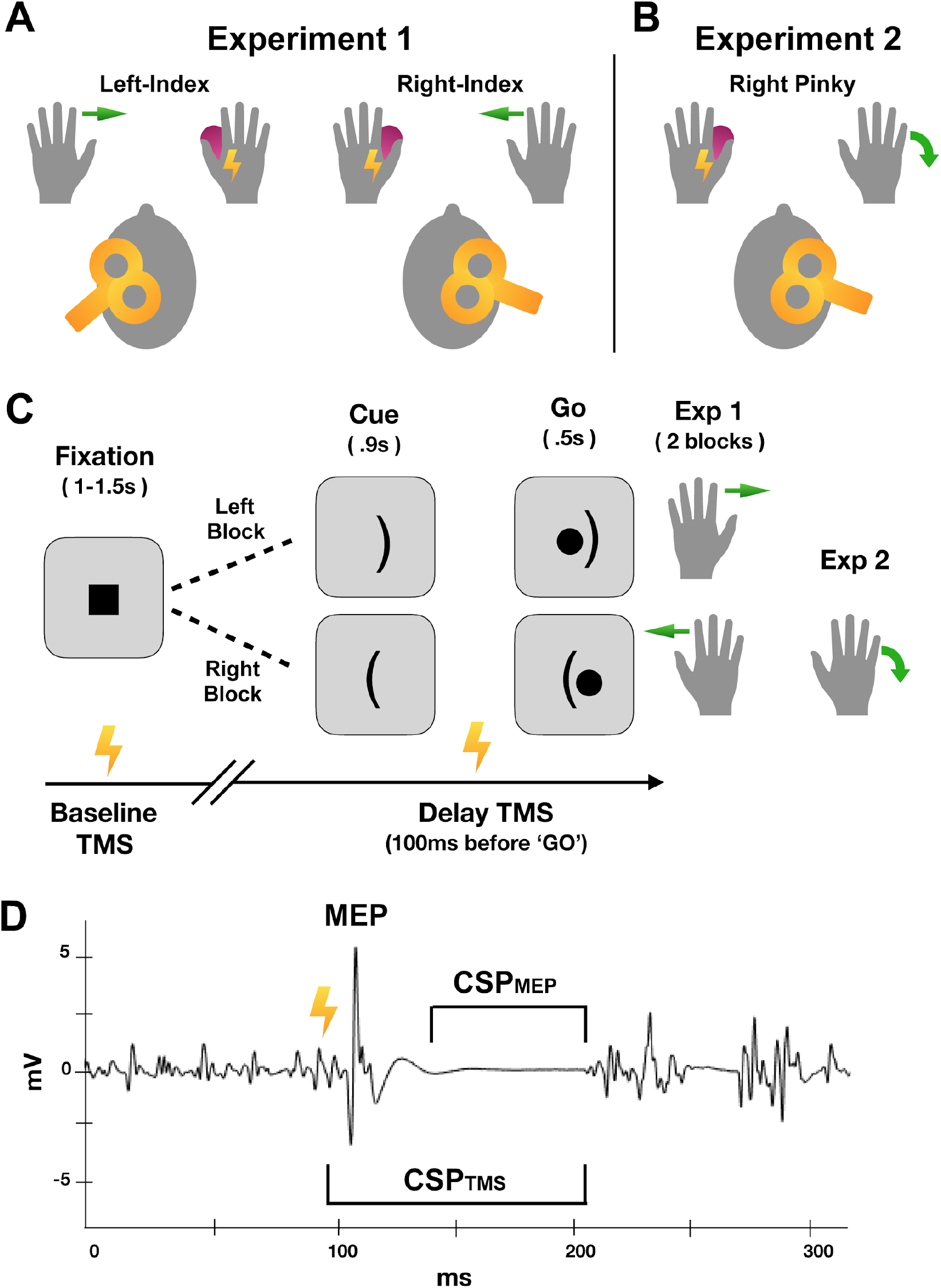
Left-index and right-index response conditions relative to TMS coil placement in Experiment 1 (A). Right pinky response condition relative to TMS coil placement in Experiment 2 (B). Timing of visual stimuli and TMS administration for the delayed response task (C). Example EMG trace showing the MEP, and the CSP measured from the TMS artifact (CSP_TMS_) and from MEP offset (*CSP_MEP_*) (D).

Tonic contraction at 25% MVC was maintained in the non-responding hand throughout each experimental block of the task, and the live EMG traces were monitored by the experimenter on an adjacent display. Verbal feedback was provided to participants, by the experimenter, if the EMG associated with the tonic contraction was outside the 25% MVC range. Participants were successful at maintaining this level of contraction for the duration of the experimental block.

TMS was delivered on 32 of 50 trials, either at the onset of the fixation cue (baseline) or 100ms prior to the imperative Go stimulus (delay), and at only one time point on a given trial. The trial order was randomized so that participants could not predict the administration or timing of TMS or whether the trial was a Go or Catch trial.

### Experiment 1

In Experiment 1, participants completed two task blocks, one with each hand, within a single testing session (Figure 1A). During one block, participants responded to the imperative stimulus by making a lateral abduction with the left-index finger to depress the response button, and the 25% MVC contraction was maintained in the right hand. In the other block, the setup was reversed, such that participants responded to the imperative stimulus by making a lateral abduction with the right-index finger to depress the response button, and the 25% MVC contraction was maintained in the left hand. The block order was counterbalanced across participants. TMS was always delivered to the primary motor cortex (M1) contralateral to the non-responding, tonically contracted hand, yielding MEP and CSP measurements from the task-irrelevant FDI muscle. With this setup, the non-responding FDI targeted by TMS was *homologous* to the responding muscle.

### Experiment 2

Experiment 2 consisted of a single task block in which the right abductor digiti minimi (ADM) was the responding muscle and the left FDI was the non-responding muscle (Figure 1B). Participants made downward pinky movements (towards the table) to depress a button on a custom-built response device designed for the right hand. As in Experiment 1, the non-responding left hand maintained a tonic contraction near 25% MVC, and TMS was administered over the right M1 to elicit MEP and CSP measurements from the task-irrelevant left FDI. In contrast to Experiment 1, this setup represents the *non-homologous* case, in which the responding muscle (right ADM) is contralateral but *non-homologous* to the tonically contracted left FDI muscle targeted by TMS.

### Data Analysis

Offline analysis of EMG data was performed using the VETA toolbox and custom automated procedures within MATLAB. Dependent variables of interest included CSP duration, MEP duration, MEP peak-to-peak amplitude, button press RT, EMG burst onset RT, and the percentage of failed catch trials. MEP duration was estimated using the Matlab *findchangepts.m* function in the window from 18 ms to 100 ms following the TMS artifact. The first point was identified as the MEP onset, and the last point as the MEP offset (Jackson & Greenhouse, 2019). CSP duration was estimated using two different approaches differing only in the identified onset of the CSP (Figure 1D). For the first approach, the CSP was estimated as the period from the TMS artifact through the resurgence of EMG activity, as in many previous studies. We refer to this estimate of the CSP duration as the CSP_TMS_ since it begins with the TMS artifact. For the second approach, the CSP was estimated as the period of MEP offset through the resurgence of EMG activity. We refer to this estimate of the CSP duration as the CSP_MEP_. The latter approach accounts for the possibility that the CSP may not be measurable until after the MEP has resolved and depends on the calculation of the MEP duration. Delay period CSP_TMS_, CSP_MEP_, MEP duration, and MEP amplitudes were calculated as a percentage of the respective baseline measurements. Button press and EMG burst onset RTs were calculated separately for no TMS, baseline TMS, and delay period TMS trials.

Two-tailed statistical tests were used to address the competing hypotheses that M1 intracortical inhibition either increases or decreases in a widespread manner during response preparation. Paired two-tailed *t*-tests compared CSP duration between the task delay period and task baseline. To evaluate possible laterality differences, we used a paired *t*-test to compare CSP duration between the left and right response blocks. Independent samples *t*-tests were used to compare CSP modulation between the homologous (Experiment 1) and non-homologous (Experiment 2) conditions. Parallel statistical tests were conducted to evaluate changes in MEP amplitude and duration. Additional paired *t*-tests compared button press and EMG onset RTs between response conditions in Experiment 1, and independent samples *t-*tests compared button press and EMG onset RTs between experiments. Pearson correlation coefficients were also calculated to assess relationships between changes in CSP duration and RT across participants to explore whether larger changes in CSP duration during response preparation correlated with faster response times.

## Results

We present the results of both experiments side-by-side to facilitate comparisons.

### CSP_TMS_ Duration

Baseline CSP_TMS_ duration differed between left-index (135 ± 38 ms) and right-index (173 ± 32 ms) response conditions (*t*(12) = 8.49, *p* < .01, *d* = 2.35), and between right-index (Exp. 1) and right-pinky response conditions (Exp. 2, 139 ± 34 ms; *t*(25,1) = 2.64, *p* < .05, *d* = 1.02; Figure 2A). The between experiment comparison between left-index (Exp. 1) and right-pinky (Exp. 2) conditions did not reach significance (*t*(25,1) = .294, *p* = .77). Thus, CSP_TMS_ duration at baseline was longer for the right-index response condition than the other two conditions. This was despite the fact CSPs were measured from the left FDI in both the right-index (Exp. 1) and right pinky (Exp. 2) response conditions, suggesting a potential effect of homology at baseline.

**Figure 2.**
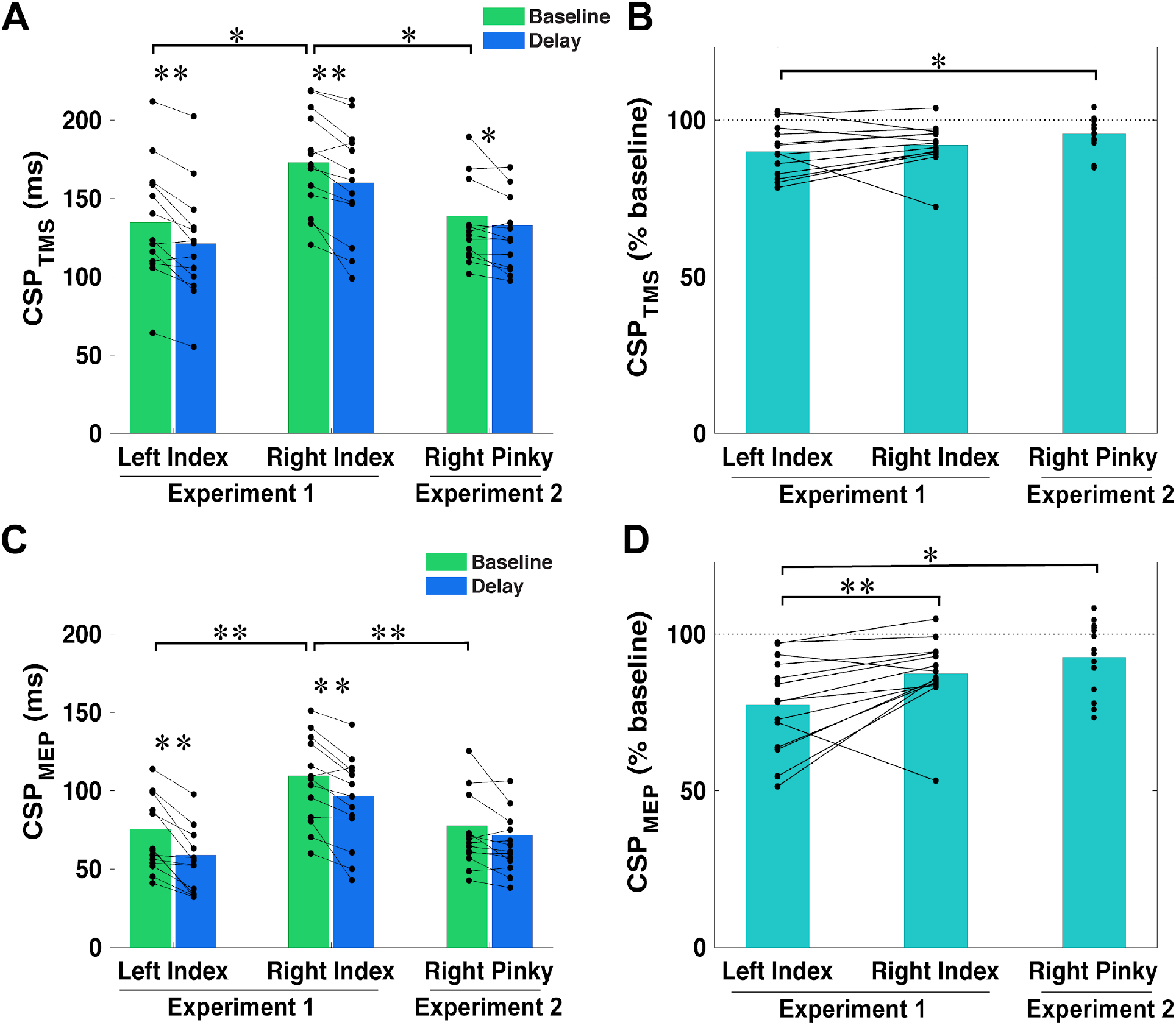
Cortical silent period (CSP) duration was significantly shorter during the preparatory delay period relative to baseline when measured from the time of the TMS pulse (A) and this decrease was larger in the left-index than right pinky response conditions (B). The pattern was similar when measuring the initiation of the CSP from the end of the MEP (C and D). * *p*<.05; ** *p*<.01.

**Figure 3.**
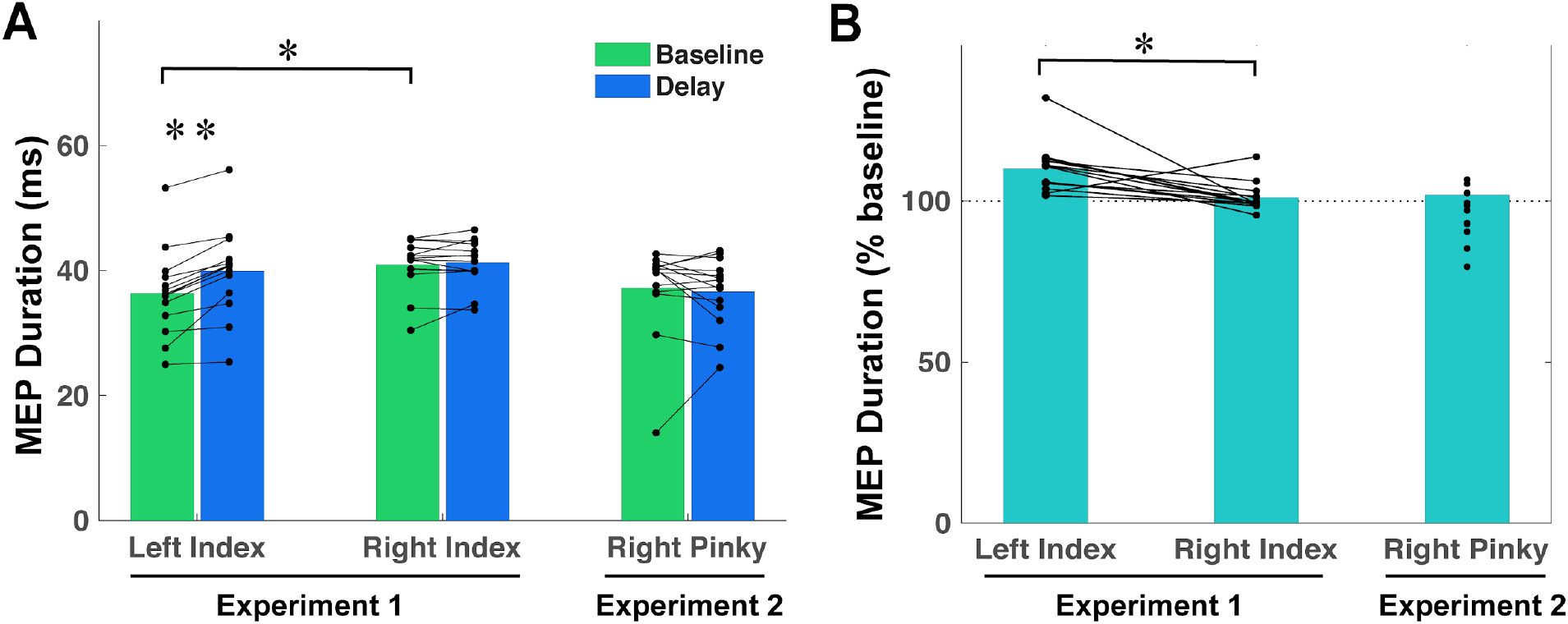
MEP duration at baseline was significantly greater for right-index compared to left-index response conditions (A). MEP duration in the left-index response condition was significantly increased during the preparatory delay period relative to baseline, and this increase was greater for the left than the right-index response condition (B). * *p*<.05; ** *p*<.01.

CSP_TMS_ duration during the preparatory delay period significantly decreased from baseline in all conditions. This was observed for the left-index (89.9 ± 8%; *t*(12) = 4.5, *p* < .01, *d* = 1.25), right-index (92 ± 7%; *t*(12) = 4.46, *p* < .01, *d* = 1.24) and right-pinky (96 ± 5%; *t*(13) = 2.73, *p* < .05, *d* = .73) response conditions (Figure 2B). The magnitude of the CSP_TMS_ decrease was significantly greater for the left-index than the right-pinky (*t*(13) = 2.19, *p* < .05, *d* = .85; Figure 2B) response condition but did not differ between the right and left-index (*t*(13) = .9, *p* = .34) or right-index and right pinky (*t*(25,1) = 1.5, *p* = .14) conditions.

To check whether CSP changes could have arisen from changes in the background tonic EMG during the response period, we analyzed the RMS of the tonic EMG signal in the 100 ms epoch following the EMG burst onset in the opposite hand (response epoch) and compared it with the 100 ms epoch preceding the TMS during the delay period. The RMS EMG in the right FDI during the left-index response epoch (.23 ± .16 mV) did not increase relative to the delay epoch (.22 ± .13 mV; *p* = .87). However, the RMS EMG in the left FDI was significantly increased during the right-index response (.21 ± .12 mV) relative to the delay epoch (.17 ± .09 mV; *t*(13) = 4.1, *p* < .01, *d* = 1.08). A similar pattern was observed for the right-pinky response condition which showed a significant increase in RMS EMG during the response epoch (.11 ± .05 mV) relative to the delay epoch (.09 ± .04 mV) (*t*(13) = 3.6, *p* < .01, *d* = 0.96). Thus, while the pattern of CSP durations did not differ across response conditions, there was a laterality difference in background EMG activity from the delay period to the response.

### CSP_MEP_ Duration

The pattern for CSP_MEP_ duration was similar to that observed for CSP_TMS_ duration, suggesting differences in the estimated MEP duration did not greatly influence CSP estimates. Baseline CSP_MEP_ duration was longer for right-index (109 ± 30 ms) than left-index (75 ± 29 ms) response conditions (*t*(13) = 6.0, *p* < .01, *d* = 1.60), and for right-index (Exp. 1) than right-pinky conditions (Exp. 2; 78 ± 30 ms; *t*(26,1) = 2.85, *p* < .01, *d* = 1.02; Figure 1C). The comparison between left-index (Exp. 1) and right-pinky (Exp. 2) did not reach significance (*t*(26,1) = .19, *p* = .85)

CSP_MEP_ duration during the preparatory delay period significantly decreased from baseline for the left-index (77 ± 15%; *t*(13) = 5.49, *p* < .01, *d* = 1.47) and right-index (87 ± 12%; *t*(13) = 4.66, *p* < .01, *d* = 1.24) conditions, but did not reach significance for the right-pinky response condition (93 ± 11%; *t*(13) = 2.1, *p* = .06; Figure 1D). The magnitude of the CSP_MEP_ decrease was significantly greater for the left-index when compared to right-index (*t*(13) = 2.76, *p* < .05, *d* = .74) and right-pinky (*t*(26,1) = 3.03, *p* < .01, *d* = 1.16) response conditions but did not differ between the right-index and right pinky (*t*(26,1) = 1.2, *p* = .24) conditions.

### MEP Duration

Baseline MEP duration was significantly shorter for the left-index (36 ± 7 ms) than the right-index (41 ± 4 ms) response condition (*t*(13) = 2.64, *p* < .05, *d* = .7; Figure 2A), but neither condition in Exp. 1 differed significantly from the right-pinky condition in Exp. 2 (37 ± 7ms; *p*’s > .05). MEP duration during the delay period significantly increased from baseline for the left-index condition (110 ± 8%, *t*(13) = 6.13, *p* < .01, *d* =1.64), but this difference was not observed for the right-index (101 ± 22%, *p* = .4) and right-pinky (102 ± 22%, *p* = .6) conditions. The increase from baseline MEP duration in the left-index response condition was significantly greater than in right-index response condition (*t*(13) = 3.52, *p* < .01, *d* = .94), but comparisons with the right pinky response condition (*t*(13,1) = 1.3, *p* = 0.2) did not reach significance.

### MEP Amplitudes

Baseline MEP amplitudes were significantly smaller in the right pinky (4.2 ± 2.1 mV) than the left-index (6.2 ± 2.5 mV, *t*(13) = 2.24, *p* < .05, *d* = .85) and right-index (6.8 ± 2.3 mV, *t*(13) = 3.16, *p* < .01, *d* = 2.17) response conditions, possibly reflecting differences between subjects or experiments (Figure 4A). However, delay period MEP amplitudes did not differ significantly from baseline for left-index (99 ± 8%), right-index (98 ± 11%) and right-pinky (93 ± 11%) conditions, and this pattern did not differ between response conditions (Figure 4B).

**Figure 4.**
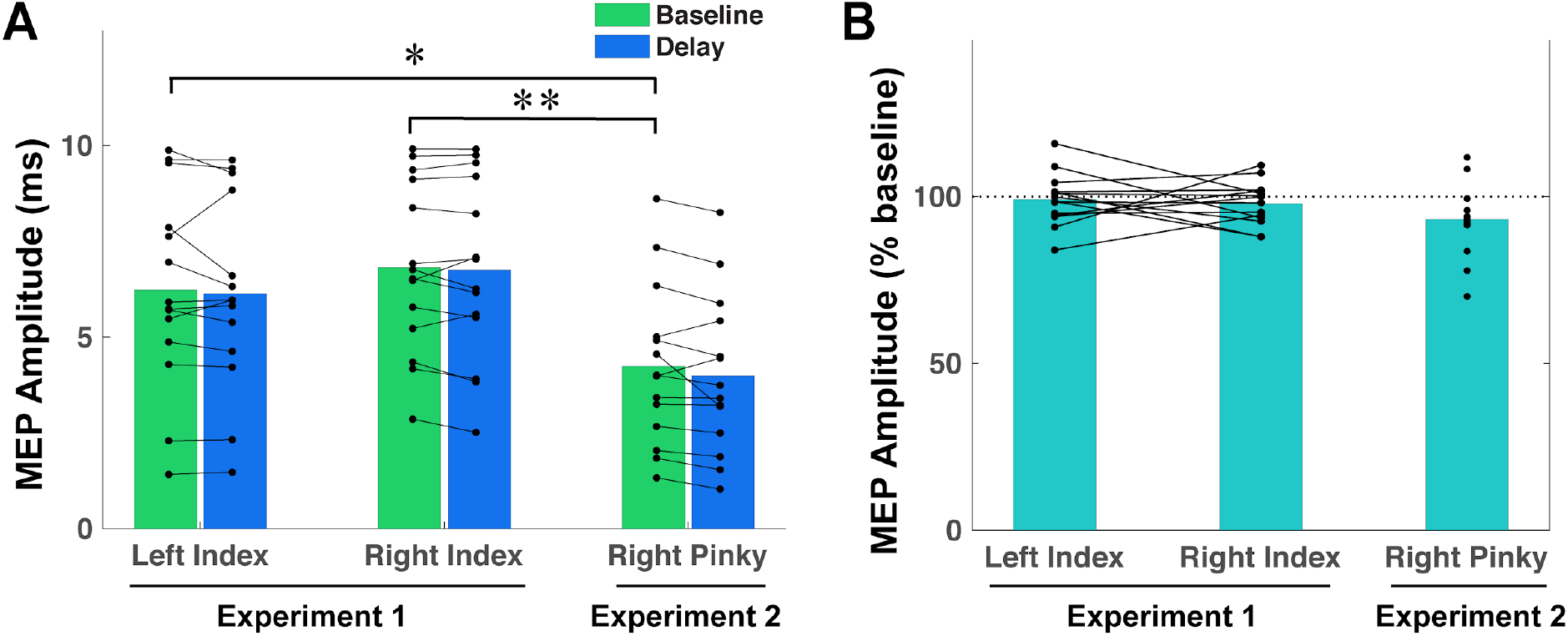
MEP amplitudes did not differ between baseline and the preparatory delay period (A) and did not differ across response conditions. MEP amplitudes were greater in Experiment 1 than Experiment 2. * *p*<.05; ** *p*<.01.

**Figure 5.**
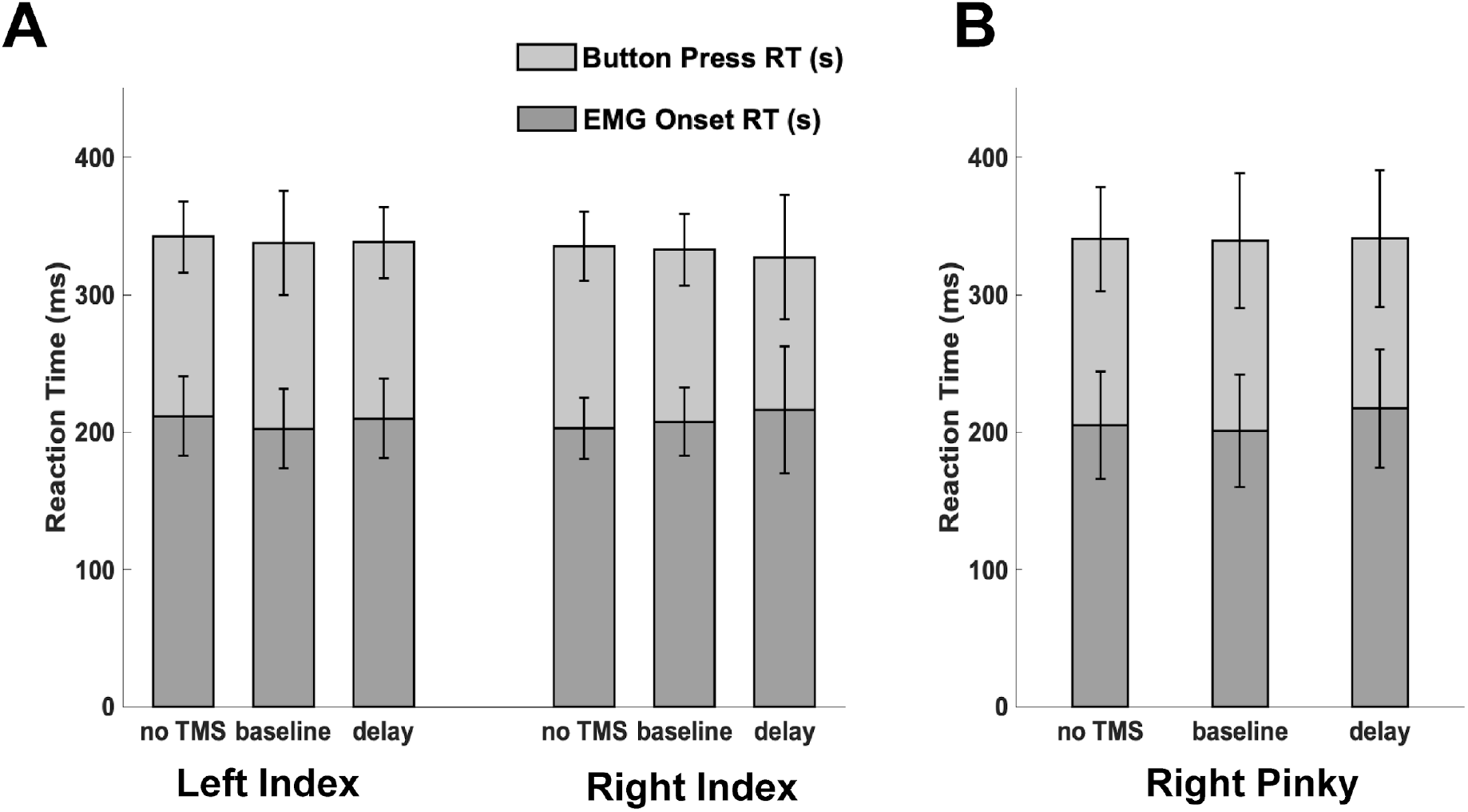
Button press reaction times (RTs) and EMG burst onset RTs did not show differences across hands or TMS conditions in Experiment 1 (A) or across TMS conditions in Experiment 2 (B). There were no differences between Experiments 1 and 2 as well.

### Overall and Pre-TMS tonic EMG

The maximum tonic EMG in the nonresponding hand averaged across all trials was 1.35 ± 0.71 mV in the left-index response condition in Experiment 1, 1.34 ± 0.81 mV in the right-index response condition in Experiment 1, and 0.73 ± 0.30 mV for Experiment 2. RMS of the EMG signal in the 100 ms preceding TMS for the left-index response condition was not significantly different between baseline (0.22 ± 0.14 mV) and delay periods (0.22 ± 0.13 mV; *p* = .66). However, there were significant differences between the baseline and delay periods for the right-index response condition (baseline: 0.19 ± 0.12 mV and delay: 0.17 ± 0.1 mV; *t*(13) = 2.44, *p* < .05, *d* = .65) and right-pinky condition (baseline: 0.10 ± 0.04 mV and delay: 0.09 ± 0.04 mV; *t*(13) = 2.69, *p* < .05, *d* = .72). Therefore, in the left-index response condition, background EMG differences did not account for the observed differences in the CSPs or MEPs. In the other two cases, the pattern indicates a decrease in background EMG activity during the preparatory period, which one might expect to correspond to an increase rather than a decrease in CSP duration.

### Button press and EMG burst onset RTs

Button press RTs for Experiment 1 did not differ between left-index (noTMS: 342 ± 26 ms; baselineTMS: 338 ± 26 ms; delayTMS: 338 ± 26ms) and right-index (noTMS: 335 ± 25 ms; baselineTMS: 332 ± 26 ms; delayTMS: 327 ± 45ms) responses, and there was no effect of TMS or TMS timing (all *p*’s > . 05). Right pinky response RTs in Experiment 2 (noTMS: 341 ± 38 ms; baselineTMS: 340 ± 49 ms; delayTMS: 341 ± 50 ms) did not differ from left-index or right-index responses in Experiment 1 (all *p*’s > .05). There was also no effect of TMS in Experiment 2 (all *p*’s > .05).

A similar pattern was found for EMG onset RTs. There was no effect of response finger in Experiment 1 (*p* > .05), no effect of TMS for either experiment (all *p*’s > .05), and no difference in EMG burst onset RTs between experiments (all *p*’s > .05). Thus, the CSP and MEP results were not explained by differences in response performance.

## Discussion

In this study, we tested the competing hypotheses that intracortical inhibition increases or decreases in task-irrelevant muscles during the preparation of actions. We found compelling evidence for a decrease in intracortical inhibition during action preparation. Specifically, we observed reductions in the CSP duration measured in a task-irrelevant muscle whether it was homologous (Experiment 1) or non-homologous (Experiment 2) to the contralateral responding muscle. This pattern was not explained by MEP amplitude, MEP duration, or background EMG activity levels. The observed changes in CSP duration in the absence of changes in MEP amplitudes implicate a mechanism involved in the cortical release of GABA-ergic inhibition distinct from mechanisms that influence properties of the MEP.

### Interpretation of reduced intracortical inhibition

Several studies have characterized the spinal and cortical contributions to the CSP. Seminal work attributed the first 50-80 ms of the CSP to a spinal origin (Fuhr et al., 1991; Ziemann et al., 1993), and the later portion to the interruption of voluntary cortical drive (Fuhr et al., 1991). These early findings would later be supported by epidural recordings (Chen et al., 1999) and pharmacological studies (Pierantozzi et al., 2004; Siebner et al., 1998; Werhahn et al., 1999; Ziemann et al., 1996). While the spinal contribution remains relatively stable, the cortical contribution is thought to reflect GABA-ergic intracortical mechanisms, and be primarily responsible for changes in the CSP duration (Siebner et al., 1998; Werhahn et al., 1999). Whether such effects are specific to GABA_A_ or GABA_B_ mediated mechanisms remains the subject of debate (McDonnell et al., 2006; Paulus et al., 2008). Nevertheless, given the existing evidence for sources of the CSP, we believe our results are best explained by a release of inhibition at the cortical level.

Short-interval intracortical inhibition (SICI) is another TMS-derived index of cortical inhibition which manifests as a reduction in the MEP amplitude when a conditioning TMS pulse is administered 1-5 ms prior to a MEP eliciting test pulse (Kujirai et al., 1993) and is widely accepted to reflect fast-acting GABAA – dependent inhibition (Lazzaro et al., 2005; Lazzaro et al., 2006). SICI is decreased during action preparation, consistent with a transient release of fast intracortical inhibition (Duque & Ivry, 2009). SICI Similarly, long-interval intracortical inhibition (LICI) manifests as a reduction in MEP amplitude when a suprathreshold conditioning stimulus precedes a test stimulus by ~100-200 ms (Nakamura et al., 1997; Valls-Solé et al., 1992). In a warned reaction time task, SICI and LICI in the responding muscle were reduced during a short preparatory fore-period, indicating a release of inhibition (Sinclair & Hammond, 2008). These results suggest intracortical inhibition is reduced during action preparation in task-relevant muscles. However, task-irrelevant muscles were not investigated in these studies.

In the context of this previous work, our findings suggest there is a release of intracortical inhibition during action preparation involving multiple GABA-dependent mechanisms acting across different time scales and with potentially different spatial distributions. Our data show that the release of intracortical inhibition in the form of reduced CSP duration, previously observed in task-relevant (Davranche et al., 2007) muscles, extends to task-irrelevant muscles as well. Further studies should investigate whether SICI and LICI change in task-irrelevant muscles during response preparation to address the question of whether the spatial extent of the release of intracortical inhibition is shared across multiple mechanisms or could dissociate between different cortical inhibitory mechanisms.

### Motor evoked potentials and laterality differences

MEP duration also showed an interesting pattern, exhibiting a significant increase from baseline to delay, but only for left-index response trials. We also note there were marked laterality differences in CSP duration. These likely reflect the influence of hand-dominance, as CSPs were shorter in the dominant hand. This finding replicates previous work which reported hand-dominance effects on CSP without changes in MEP amplitude, latency, and threshold (Priori et al., 1999). MEP amplitudes did not exhibit a similar pattern, pointing to an interesting dissociation between MEP duration and MEP amplitude.

Interestingly, we observed a decrease in the RMS of the EMG activity immediately preceding the TMS pulse between baseline and delay period measurements, but this was only the case for right hand responses, i.e. when tonic EMG was recorded from the left FDI. We also observed increased tonic EMG activity during the transition from the delay period into the response phase but only for right hand responses as well. These patterns may reflect laterality differences, although opposite the pattern we observed for MEP duration. Moreover, this did not appear to influence MEP amplitudes.

While hand-dominance may explain the differences between left-index and right-index responses, we were unable to compare the differences between the index and pinky response conditions within participants. Therefore, differences between the right-index and right pinky response conditions may be explained by inter-subject differences.

### Alternative interpretations and limitations

The CSPs and MEPs were measured from tonically active muscles, which complicates comparisons of our results to previous investigations of preparatory inhibition. We did not observe a decrease in MEP amplitudes during response preparation, a finding from multiple previous studies (Duque et al., 2010; Duque & Ivry, 2009; Greenhouse et al., 2015; Hasbroucq et al., 1997; Ibáñez et al., 2020; Labruna et al., 2019; Lebon et al., 2016; Sinclair & Hammond, 2008; and others). This could be explained by the level of tonic EMG maintained in the muscle from which CSPs were measured. By asking participants to remain in a tonically active state, we may have washed out the commonly observed preparatory MEP suppression and, in exchange, uncovered a decrease in CSP duration. Previously, authors measured MEPs in task-irrelevant muscles *while at rest*, which deliberately avoids the possible interference introduced by tonic EMG activity. We note that one previous study observed reduced MEP amplitudes during response preparation in a tonically active muscle (Hannah et al., 2018) although the muscle was task-relevant and tonic contraction was 5-10% MVC, a lower intensity than we used here. In contrast, the design of the current study, which required participants to maintain 25% MVC, likely introduced interhemispheric effects and stronger descending corticospinal drive. Interestingly, previous work suggests a low-intensity but not a high-intensity contraction results in decreased MEP amplitudes in a task-irrelevant homologous muscle (Liepert et al., 2001).

We also note that RMT served as the reference for determining the TMS intensity used for CSP measurements, rather than an active motor threshold (AMT) as has been frequently used in other studies. This yielded MEP amplitudes larger than those found in most previous studies. The higher level of activation may have further impacted our ability to detect changes in MEPs during the task. On a separate note, even with these large MEP amplitudes, we did not observe a difference in reaction times between trials with and without TMS, raising questions about the possible source of RT differences reported in previous work. In the response hand, this pattern of faster RTs in the presence of TMS may reflect the influence of a shortened CSP.

Since we only measured EMG from hand muscles, we cannot make strong claims about the widespread nature of CSP modulation in other task-irrelevant muscles. Similarly, it is unclear whether the CSP results from GABAA and GABAB mechanisms. Comparing SICI, LICI, and CSP measurements will help to elucidate which of these mechanisms may be responsible for preparatory modulation in task-irrelevant muscles.

### Conclusion

We observed evidence of a nonfocal release of intracortical inhibition during the preparation of actions evident in the form of decreased duration of CSPs measured from task-irrelevant muscles. This pattern was detected in the absence of a change in MEP amplitudes. Our findings are consistent with a model in which response preparation involves a widespread release of cortical inhibition and extend those of previous studies that reported changes in other TMS-derived markers of intracortical inhibition in muscles involved in the task. Furthermore, given that we did not observe concurrent changes in MEP amplitudes, our results suggest the release of intracortical inhibition arises from a mechanism that operates independent of other mechanisms that influence corticospinal excitability. These findings have clinical relevance for diseases which impair response initiation, such as Parkinson’s disease, stroke, and trauma by providing insight into potentially affected mechanisms.

## Notes

https://osf.io/jvnmq/

### Competing Interest Statement

The authors have declared no competing interest.

